# Combining *in vivo* 2-photon imaging with photoactivatable fluorescent labeling reveals strikingly low rates of mitochondrial dynamics in skeletal muscle

**DOI:** 10.1101/2024.12.10.627796

**Authors:** Colleen L. O’Reilly, Arik Davidyan, Katarzyna Cizio, Stephen M. Doidge, Matthew P. Bubak, Agnieszka K. Borowik, Tommy L. Lewis, Benjamin F. Miller

**Affiliations:** Aging and Metabolism Research Program, Oklahoma Medical Research Foundation, Oklahoma City, Oklahoma, USA; Department of Biological Sciences, Sacramento State University, Sacramento, California, USA; Neuroscience Program, University of Oklahoma Health Sciences Center, Oklahoma City, Oklahoma, USA; Oklahoma City Veterans Association, Oklahoma City, OK, USA

**Keywords:** Mitochondrial Dynamics, Fusion, Fission, Skeletal Muscle

## Abstract

Mitochondrial dynamics involve two distinct and opposing processes, fusion and fission. Traditionally we assess fusion and fission by snapshots of protein markers at distinct time points or *in vitro* models to infer outcomes *in vivo*. Recent technological advancements enable visualization of mitochondrial dynamics *in vivo* using fluorescent microscopy. Our study modified this technique to evaluate mitochondrial dynamics in skeletal muscle, comparing young (6mo) and old (24mo) mice *in vivo* and contrasting this to *ex vivo* and *in vitro* models. We hypothesized that *in vitro* and *ex vivo* models would have higher rates of dynamics than *in vivo* models and that young animals would have higher rates than old animals. We electroporated mitochondrial matrix-targeted photo-activatable GFP into the tibialis anterior (TA) of young and old C57Bl6 mice and imaged using multiphoton microscopy. We also measured rates of mitochondrial dynamics using single fibers isolated from the TA of the electroporated mice, as well as C2C12 myotubes transfected with the same plasmids. We found that the rates of dynamic events *in vivo* are slower than previously indicated, with the C2C12 myoblasts having the fastest rates of dynamic events across all models. We also observed that dynamic rates are slower in old animals compared to young animals. Finally, we found that rates of dynamic events were higher in old animals after an acute bout of exercise. Our data demonstrate it is possible to directly measure rates of mitochondrial dynamics *in vivo*. This technique provides a powerful tool to answer experimental questions about mitochondrial dynamics of skeletal muscle.

## Introduction

Skeletal muscle is the largest organ in the body by mass and plays a critical role in mobility, protein homeostasis, and energy metabolism. With age, there is a loss of muscle mass and strength, associated with functional impairment, physical disabilities, and an overall decline in quality of life (1, 2). Sarcopenia is a driving force in the aging process and the healthcare burden of aging (3). Several different mechanisms have been proposed to explain the development of sarcopenia (4), but evidence suggests that several components of mitochondrial function play a role in muscle aging processes (5–7).

Skeletal muscle mitochondria form an intricate, interconnected network that spans the entire length and depth of muscle fibers (8–12) in order to distribute energy throughout the cell (13). Two important remodeling processes govern the structural organization of mitochondria in skeletal muscles: protein turnover and mitochondrial dynamics. In skeletal muscle, mitochondrial dynamics involves two distinct and opposing processes termed fusion and fission. Fusion is the process of two mitochondria merging into one larger mitochondrion, while fission involves one mitochondrion breaking into multiple smaller mitochondria. Fusion is principally controlled by three mitochondrial GTPases: inner membrane-localized optic atrophy-1 (OPA1) (14) and two outer membrane-localized mitofusins (MFN1 and MFN2) (15). These three proteins are often used as markers of fusion (16–18). Fission is controlled by several proteins, including dynamin-related protein-1 (DRP1), mitochondrial fission factor (MFF), fission-1 (FIS1), and mitochondrial elongation factors 1 & 2 (MIEF1 & 2). The recruitment and association of DRP1 to the mitochondria is mainly driven by MFF and FIS1 while phosphorylation of DRP1 at Ser616 stimulates DRP1 oligomerization and activates mitochondrial fission (19). The use of preclinical animal models in which fusion and fission are genetically manipulated shed light on the critical nature of fusion and fission processes for maintaining mitochondrial, and thus skeletal muscle function (Reviewed in citation 20). However, there is still little consensus on how changes in mitochondrial dynamics impact skeletal muscle aging.

Methods that capture the dynamic nature of mitochondrial protein turnover highlight the limitations of using protein markers to understand remodeling events (21). Although there are good methods for measuring protein remodeling, assessing the role that mitochondrial dynamics play in both health and disease states of skeletal muscle is still hindered by a reliance on protein markers, *in vitro* assays, and single time point assessments. In neurons (22), the sciatic nerve (23), and skin (24), methods have been developed to directly assess mitochondrial dynamics *in vivo* to avoid reliance on indirect measures. *In vivo* analysis of mitochondrial dynamics in cortical neurons suggests that increased fission via MFF activity determines the size of axonal mitochondria (25). Additionally, *in vivo* analysis of mitochondria within epithelial layers using NADH two photon excited fluorescence revealed a distinct temporal difference between mitochondrial biochemistry and dynamic morphology, with fusion having a larger delay between the two than fission (24). Despite these advances in other tissues, researchers have yet to establish an approach for quantifying rates of mitochondrial dynamics i*n vivo* for skeletal muscle.

Our current understanding of skeletal muscle mitochondrial dynamics has progressed mainly due to improved imaging and labeling techniques. For example, imaging of primary cultures of flexor digitorum brevis myofibers suggests that mitochondrial fusion events occur in skeletal muscle fibers at rates of nearly 0.5 events/minute (26). Additionally, work in *ex vivo* extensor digitorum longus (EDL) muscle indicate that mitochondrial dynamics is tailored to the specific metabolic state of different skeletal muscle fiber types (27). However, investigations of mitochondrial remodeling *in vitro* likely do not represent the conditions *in vivo.* For example, protein synthesis rates *in vitro* (28, 29) are higher than those measured *in vivo* (21, 30). These differences between *in vivo* and *ex vivo* are most likely because *in vitro c*onditions are substrate-dense and lack the complexity of *in vivo* environments. A study using *in vivo* imaging at a single time point showed that inhibition of mitochondrial fission prevents fasting-induced muscle atrophy (31). However, like when using a marker, a single time point cannot capture changes in the rates of mitochondrial dynamic events and therefore may lead to erroneous conclusions. Finally, there was one study that combined electroporation of a photoactivatable fluorescent tag and confocal microscopy to investigate changes in mitochondrial dynamics over time in muscle tissue in a rodent model of type II diabetes (32). However, factors such as the time period assessed, and the potential pitfalls of short wavelength laser-induced phototoxicity (33) could have all impacted the measured rates of mitochondrial dynamics. Although this paper performed *in vivo* imaging, it did not report rates of the dynamic events. Quantifiable rates of mitochondrial dynamics provide a better understanding of the remodeling processes (fusion and fission) independent of observable morphological changes. Improving the ability to monitor and measure quantifiable rates of mitochondrial dynamics *in vivo* is an important step toward understanding its role in health, disease, aging and implementing interventions.

Our study aimed to develop and validate a technique using multiphoton microscopy to assess *in vivo* rates of mitochondrial dynamics in skeletal muscle over extended periods of time. We then compared these rates to *in vitro* and *ex vivo* models, between young and old mice, and after an acute bout of exercise. To assess rates of mitochondrial dynamics *in vivo* in skeletal muscle, we coupled electroporation of a construct encoding both a mitochondrial matrix-targeted photoactivable green fluorescent protein (mt-paGFP) and matrix targeted mScarlet (mt-mScarlet) with *in vivo* 2-photon imaging. We hypothesized that rates of fission and fusion events measured *in vivo* would be slower than those measured *in vitro* and *ex vivo*. We further hypothesized that the rates of fission and fusion events would be slower in old compared to young mice. Lastly, we hypothesized that addition of an acute bout of exercise would stimulate the rates of mitochondrial dynamics in both young and old muscle compared to sedentary controls.

## Methods

### Animals

All animal procedures were conducted in accordance with institutional guidelines for the care and use of laboratory animals and were approved by the institutional Animal Care and Use Committees at the Oklahoma Medical Research Foundation (OMRF). Young (6-month) and old (24-month) male and female C57Bl6 background mice were housed in the vivarium at OMRF. Animals were group housed (a maximum of five per cage) with *ad libitum* access to food and water in a room on 14:10 light:dark cycle with constant temperature and humidity.

### Plasmids

pCAG:2xmt-mScarlet was created by cloning a DNA gene block (IDT) encoding two CoxVIII leader sequences (2xmt) 5’ to the sequence for mScarlet into pCAG via restriction digest with Xho1 and Not1. pCAG:2x-mtpaGFP p2a 2x-mtmScarlet was created by cloning a DNA gene block (IDT) encoding 2xmt-photoactivatible GFP (paGFP) p2a 2xmtmScarlet into pCAG via restriction digest with Xho1 and Not1. pCAG:mt-GCaMP6f was created by excising the YFP from pCAG-mt-YFP (22) and inserting the sequence encoding GCaMP6f (a gift from Douglas Kim, RRID:Addgene_40755) in its place. pCAG:tdTomato was previously published in (22). All plasmids were sequenced by NGS sequencing (Plasmidsaurus) to confirm their sequence. Plasmids were prepared and purified using the Macherey-Nagel NucleoBond Xtra Midi EF kit (item number:740412.5, MACHERY-NAGEL, Duren, Germany) to ensure ultrapure, endotoxin-free plasmid preparations.

### Electroporation of Plasmids

One week prior to imaging, plasmids were electroporated into to the tibialis anterior (TA) muscle as previously described (34). Briefly, animals were anesthetized with isofluorane (4-5% induction, 1% maintenance) on an IR heating pad. We shaved the leg and applied Nair hair removal cream for 30 seconds to a minute and then wiped clean with a wet wipe ensuring that all Nair was removed. Using an insulin syringe, we injected 31 µl of 0.4 U/ml hyaluronidase (Sigma-Aldrich, St. Louis, MO, USA) into the belly of the TA muscle from the distal end and returned the mice to their home cage for recovery. Two hours after hyaluronidase injection, a total of 10 µg of plasmid DNA, for either calcium sensing (mt-GCaMP6f & 2xmt-mScarlet at a 1:1 ratio) or the photo-activatable GFP (paGFP) plasmids (see plasmid info) were prepared in 30 µl of sterile saline and injected with the mouse under isoflurane. To electroporate the skeletal muscle, we placed a small amount of Ultrasound Transmission Gel (Aquasonic 100) on the anterior and posterior of the shaved hindlimb. We then placed seven-millimeter tweezer electrodes (Harvard Apparatus, Holliston, MA, USA) around the leg where one side was located on top of the belly of the TA and the other was on the medial head of the gastrocnemius. An electrical stimulus was applied using a BTX ECM830 electroporator (Harvard Instruments) with the following settings: a voltage of 86 V, 10 pulses, a pulse length of 20 milliseconds, a distance between electrodes of 2 mm, and 0.5 milliseconds between pulses. To allow for expression of the electroporated plasmids animals returned to their home cage until imaging.

### Exercise Protocol

Mice acclimated to the treadmill for two days by walking at 10 m/min for 10 minutes. On the third day, after walking for 10 minutes at 10 m/min, the incline was increased to 5%, and the animals ran for 60 min or until exhaustion. A brush held by the operator and an electric grid at the end of the treadmill encouraged animals to run. The test was terminated if the mouse stopped responding to encouragement continuously for five seconds. Animals were immediately anesthetized with isoflurane and prepared for *in vivo* imaging.

### Imaging

All *in vivo* images of the TA muscle were taken seven days after electroporation **(Figure 1A)** using a Nikon A1R multiphoton microscope with custom stage (Nikon Instruments Inc, New York, United States), and Coherent Chameleon Vision Ultra tunable IR laser (Coherent Corp., California, United States). Mice were anesthetized with isoflurane (4-5% induction) and then transferred to the microscope stage on top of an IR heating pad with constant isoflurane through a nose cone (1% maintenance). The electroporated TA was placed on a custom 3D printed block and secured with animal safe tape **(Figure 1B)**. To stabilize the TA muscle on the 3D printed block we employed a metal headpost (Neurotar). We used a scalpel blade to cut away the skin and expose muscle tissue within the window of the headpost, after which the muscle was kept moist with Krebs buffer. Each image **(Figure 1C)** consisted of a 35-step z-stack (1024×1024 pixels) and were acquired through a 25x 1.1 NA Nikon water objective with 3x zoom using Nikon NIS elements (RRID:SCR_002776, Nikon instruments Inc., NY, USA). For imaging mt-paGFP, mt-GCaMP6f or mt-mScarlet, the laser was tuned to 960nm, and imaging performed with 5-10% laser power through NIS elements.

**Figure 1:**
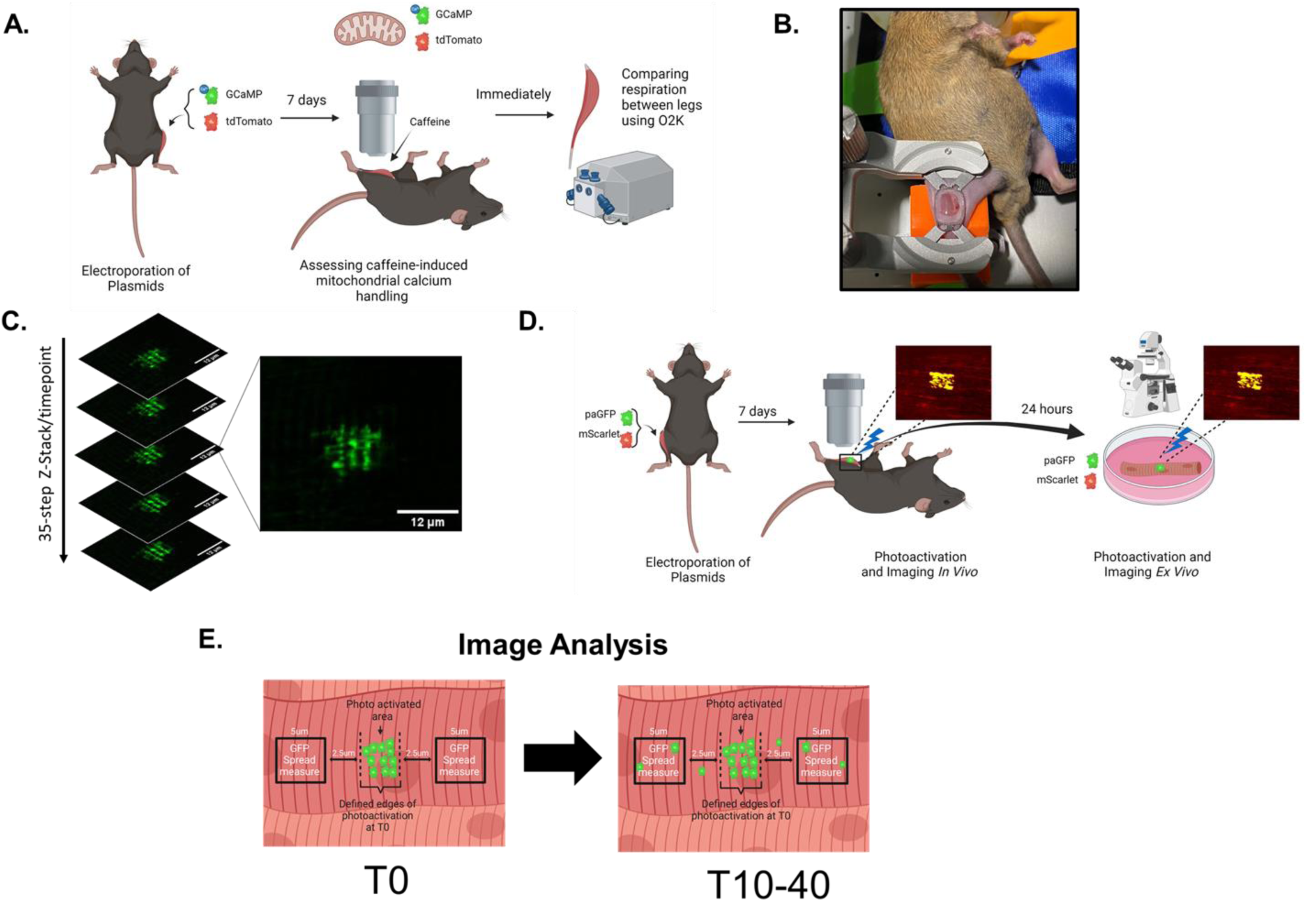
Methodology. **(A)** Methods and timeline of mitochondrial calcium uptake imaging with the 2-photon microscope and high resolution respirometry comparing the electroporated and nonelectroporated legs. **(B)** Picture of mouse mounted on 2-photon stage for live imaging of the tibialis anterior (TA). Mouse leg is mounted on a 3-D printed block and a cranial window is placed firmly across the TA muscle. Skin has been cut away and fascial layer removed. **(C)** Representative images of mt-pAGFP from one time point Z stack and single Z plane images prior to max projection **(D)** Methods and timeline of mitochondrial dynamic imaging using a photoactivatable GFP plasmid *in vivo* and *ex vivo*. **(E)** Detailed explanation of the GFP spread analysis used to measure rate of mitochondrial dynamics of max projection images across time points.

#### In vivo calcium uptake imaging

We took an image (Pre) for both the green and red channels after finding muscle fibers visually (wavelength 960 nm). The Krebs buffer bathing the muscle tissue was then removed and warm Krebs buffer with 20 mM caffeine (made fresh daily) was applied between the window and the objective. A Time 0 (T0) image was taken immediately following caffeine application, and images were taken every two minutes for up to 20 minutes. As this preparation is an intact muscle, caffeine reaches different muscle fibers across different timescales to drive mitochondrial calcium uptake over various timescales.

#### In vivo photo-activation of mt-paGFP

We took an image (Pre) after acquiring muscle fibers visually (wavelength 960nm). Regions of interest (ROI) were selected, drawn and assigned to the activation protocol. ROIs were activated using the Coherent Chameleon Vision Ultra tunable IR laser (Coherent Corp., California, United States) tuned to 760 nm, with 10 loops at 1% laser power. Based on preliminary studies demonstrating the slow rates of dynamics *in vivo*, a time zero image (T0) was taken immediately following laser activation of ROIs, with subsequent images taken every 10 minutes for 40 minutes.

#### Ex vivo photo-activation imaging

The mice were euthanized after *in vivo* imaging and the electroporated TA muscles were immediately excised. Single fibers were isolated from the excised TA muscles as previously described (35, 36). Isolated fibers were cultured in DMEM on glass bottom dishes coated with gelatin (Matek) and imaged 24 hours later with a Nikon Ti2-E microscope (Nikon Instruments Inc., New York, United States) in a stage top incubator (Okolab USA Inc., Pennsylvania, United States) (**Figure 1D)**. We took a pre-image of single muscle fibers before stimulation. Similarly to the *in vivo* imaging, ROIs were drawn and assigned to a stimulation protocol carried out with an Optimicroscan XY galvo scanning unit (405nm stimulation laser at 2% laser power for 100µs per pixel). We took a T0 image immediately after stimulation and then again, every 10 minutes for up to 40 minutes as previously described with the *in vivo* imaging.

#### In vitro photo-activation imaging

We cultured a C2C12 myogenic cell line (ATCC, Manassas Virginia) in DMEM supplemented with 10% Fetal Bovine Serum on 20 mm with 0.16-0.19 mm glass bottom petri dishes (MatTek) coated with PDL diluted to 200 µg/mL in tissue culture grade water (Sigma, St. Louis, MO, United States). Myoblasts were transiently transfected using JetPrime transfection reagent as previously described (37). We imaged the myoblasts 24 hours after transfection on a Nikon Ti2-E microscope (Nikon Instruments Inc., New York, United States), with a 60x oil immersion objective in a stage top incubator at 37° C (Okolab USA Inc., Pennsylvania, United States). Myoblasts were then differentiated into myotubes in DMEM supplemented with 10% horse serum (Gibco DMEM, 21063029). Four days after differentiation, we imaged the myotubes using the same procedure as above. The cells were stimulated in the same manner as the isolated single fibers. During imaging, myoblasts and myotubes were incubated in phenol red-free DMEM with 10% horse serum. Cells were imaged every 5 seconds for a total of 10 minutes.

### Image Analysis

We measured GFP signal intensity over time to measure calcium uptake into the mitochondria. A ROI was drawn around fibers in the image and the intensities of both green and red signals were analyzed. For each fiber ROI we created a ratio of mt-GCaMP (green) signal to cytoplasmic tdTomato (red) signal and plotted the ratio for each time point. We analyzed multiple fibers per animal (n=2-3 fibers). Each timepoint represents the average of all imaged fibers in each animal.

To assess the rates of mitochondrial dynamics, we measured the spread of GFP throughout the mitochondrial network over time **(Figure 1E)**. First, the boundaries of the initial stimulated area were determined, and then 5-micron boxes were drawn on either side of the photoactivated area. The GFP-signal intensity in those 5-micron boxes was averaged together to measure the spread of GFP at each time point. Each 5-micron box had GFP intensity corrected for bleaching via mScarlet signal relative to T0. F/F0 was calculated to determine the change in GFP intensity over time.

### High-resolution respirometry

To confirm that electroporation of the TA muscles did not impact mitochondrial function, we used high-resolution mitochondrial respiration (Oxygraph-2k, Oroboros Instruments, Innsbruck, Austria), adapted from our previously established protocols (38–40). We isolated fiber bundles from the middle third of the TA muscle and mechanically separated and chemically permeabilized with saponin (40uM). Oxidative phosphorylation and electron transport capacity were determined using the following order and concentration of substrates: NADH pathway substrates glutamate (10mM, Sigma G1626), malate (2mM, Sigma M1000), and pyruvate (5mM, Sigma P2256), adenosine 5’-diphosphate (5mM, Sigma A5285), succinate (10mM, Sigma S2378) and the uncoupler CCCP (0.75uM, C2759). Cytochrome C (10μm, Sigma C2506) was introduced before succinate to confirm membrane integrity and only runs where membrane integrity was confirmed (<10% increase in respiration after cytochrome c introduction) are presented. All presented data were normalized to respiration in the presence of antimycin A (5mM, Sigma A8674).

### Markers of mitochondrial remodeling

To assess markers of mitochondrial remodeling, we used 20–30 mg of homogenized TA muscle and isolated the mitochondrial protein fraction as previously described (41). Protein concentration was determined by Peirce660 assay (cat#: 22660, ThermoFisher Scientific, Waltham, MA, USA) following manufacturer instructions. Mitochondrial protein fractions were processed by Protein Simple Peggy Sue (San Jose, CA, USA) capillary western blot system for size-based separation and detection of the proteins according to the standard instrument protocol. The following primary antibodies were used: MFN1 (1:25, Cell Signaling Technology Danvers, MA, USA Cat# 14739, RRID:AB_2744531), MFN2 (1:10, Cell Signaling Technology Cat# 11925, RRID:AB_2750893), DRP1 (1:10, Cell Signaling Technology Cat# 8570, RRID:AB_10950498), p-DRP1 (1:25, Cell Signaling Technology Cat# 4494, RRID:AB_11178659), MFF (1:25, Cell Signaling Technology Cat# 84580, RRID:AB_2728769), OPA1 (1:25, Cell Signaling Technology Cat# 80471, RRID:AB_2734117), TOM20 (1:10, Cell Signaling Technology Cat# 42406, RRID:AB_2687663). Due to the capillary system used molecular weights may vary from traditional known values by up to 20kDa. The average molecular weight of the capillary for each antibody is included on the representative images. Secondary antibodies used for this analysis were included in the Peggy Sue Kit (SM-S001, Protein PeggySue Protein Analyzer (RRID:SCR_020406). Analyzed targets were normalized to total protein measured with the total protein detection model (DM-TP01, ProteinSimple, San Jose, CA, USA).

### Statistical Analysis

All data are presented as means ± standard deviation. For comparisons between two groups, we used unpaired t-tests. For analysis of oxygen consumption after each substrate addition in the respirometry study, we used paired t-tests to compare control and electroporated TA fiber bundles. We used a one-way ANOVA to compare the four imaging models as well as the rates of mitochondrial dynamic events in exercised animals.

## Results

To confirm that our approach was innocuous with regards to mitochondrial function, we performed mitochondrial calcium imaging followed by high-resolution respirometry. We chose to use the animals electroporated with mt-GCaMP because this plasmid binds calcium making it more likely to impact the mitochondrial respiration compared to the more inert paGFP in the subsequent experiments. Intramitochondrial calcium, measured by mt-GCaMP signal intensity normalized to cytoplasmic tdTomato signal, increased over time following caffeine stimulation **(Figure 2A-2C),** confirming calcium uptake by mitochondria. This result indicates that our method is sensitive enough and at a high enough resolution to capture both dynamic and subtle changes within the mitochondria of the intact TA muscle. Immediately following the imaging, high-resolution respirometry was completed to assess the potential effect of electroporation on mitochondrial function **(Figure 2D)**. We found no differences in mitochondrial respiration between the electroporated and nonelectroporated muscles at maximal complex 1 oxidative phosphorylation (ADP), complex 1 and 2 OXPHOS (S), and electron transport capacity (CCCP), confirming that the electroporation and subsequent imaging of the TA muscle did not alter mitochondrial respiration capacity. Coupled together, these results demonstrate that our in vivo imaging approach does not negatively impact mitochondrial function.

**Figure 2.**
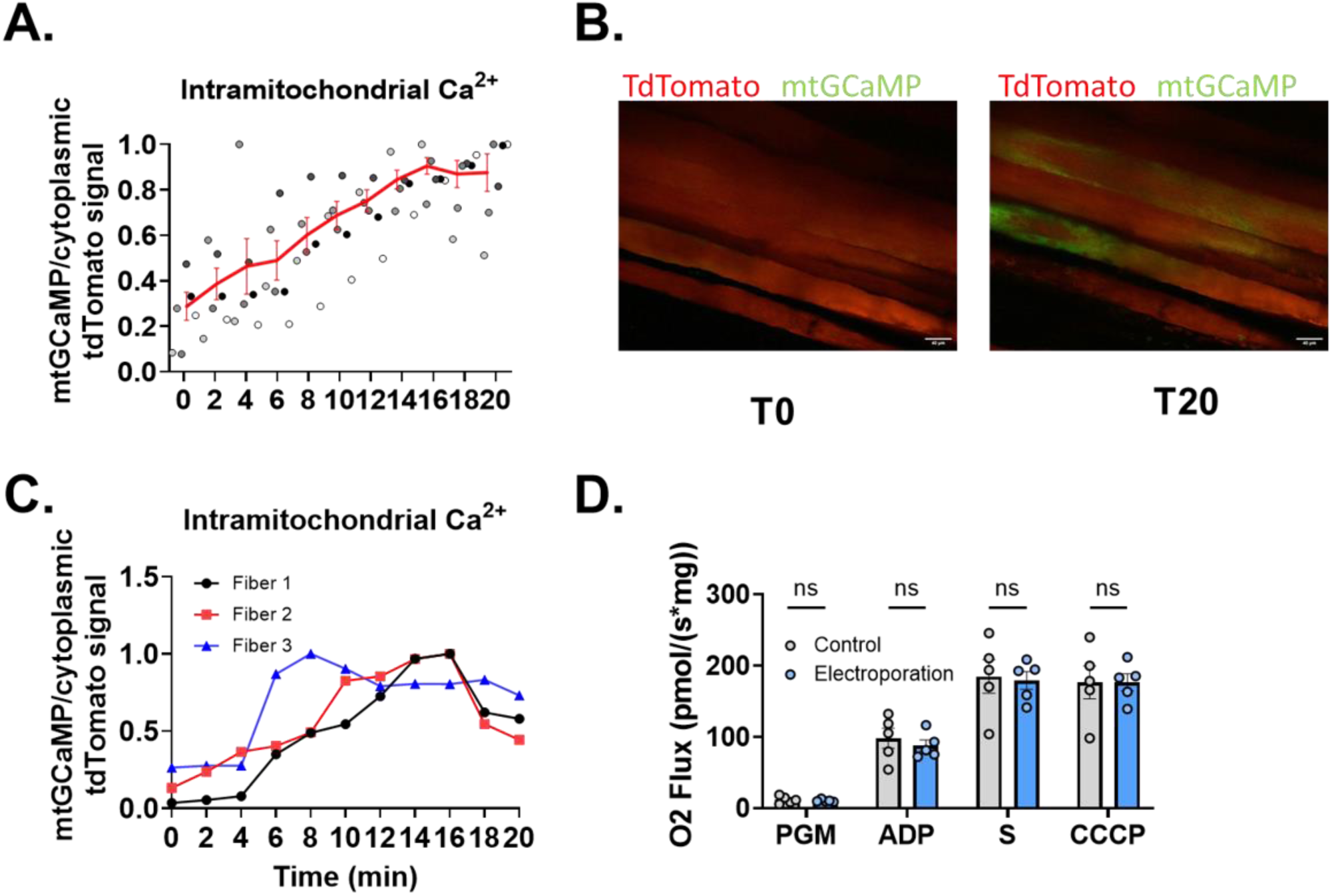
*In vivo mitochondrial GCaMP* imaging and respiration comparison. (**A)** mt-GCaMP signal normalized to cytoplasmic tdTomato signal over 20 minutes. Red line illustrates the average of all animals. For each fiber ROI we created a ratio of mt-GCaMP (green) signal to cytoplasmic tdTomato (red) signal and plotted the ratio for each time point. We analyzed multiple fibers per animal (n=2-3 fibers). Each dot represents the average of all fibers analyzed for a particular animal. **(B)** representative images of mt-GCaMP (green) and cytosolic TdTomato (red) at T0 and T20. Scale = 40µm **(C)** representative fibers showing that intramitochondrial calcium went up upon caffeine administration and then began to decline. Contrast increased in both for visual clarity. Each image consisted of a 1024×1024 pixels image and were acquired through a 25 x 1.1 NA Nikon water objective with 1x zoom using Nikon NIS elements (Nikon instruments Inc., NY, USA **(D)** O2 Flux comparisons of control and electroporated legs of the mt-GCaMP imaged animals (n=5). Paired t-tests were performed for each substrate. Data are presented as means ± SD. ns=non-significant.

Previous investigations *in vitro* and *ex vivo* suggest high rates of mitochondrial dynamic events in skeletal muscle but to our knowledge *in vivo* and *ex vivo* models have never been compared. To determine if *in vitro* and *ex vivo* approaches approximate the rates of dynamics found *in vivo,* we compared the rates of mitochondrial dynamics across experimental models. To do this we measured the GFP spread into 5 µm regions of interest (measurement ROIs) that were 2.5 µm from the original photo-activation region. We empirically determined that our reported GFP movement into these ROIs was due to fission or fusion events and not from dispersion within interconnected mitochondria (**Figure 1D**). Because there was a significant difference in the timing of GFP spread between *in vivo*, *ex vivo*, and *in vitro* conditions, we opted to make the comparison between the groups at a 10-minute time point. While the spread of GFP into measurement ROIs at 10 minutes was quite robust for both myoblasts and myotubes, there was no increase in GFP intensity between 0 and 10 minutes in the measurement ROIs for either *in vivo* or *ex vivo* fibers **(Figure 3A-D, Supplemental Figure 1).** When comparing the change in GFP spread over 10 minutes, the *in vitro* models were significantly higher than either the *in vivo* or *ex vivo* models **(Figure 3E).** Additionally, even at 40 minutes post-stimulation, we detected minimal GFP spread in both the *in vivo* **(Figure 4A)** and *ex vivo* model **(Supplemental Figure 2).** These results demonstrate that the rates of mitochondrial dynamics in intact skeletal muscle fibers are much lower than in cultured C2C12 models.

**Figure 3.**
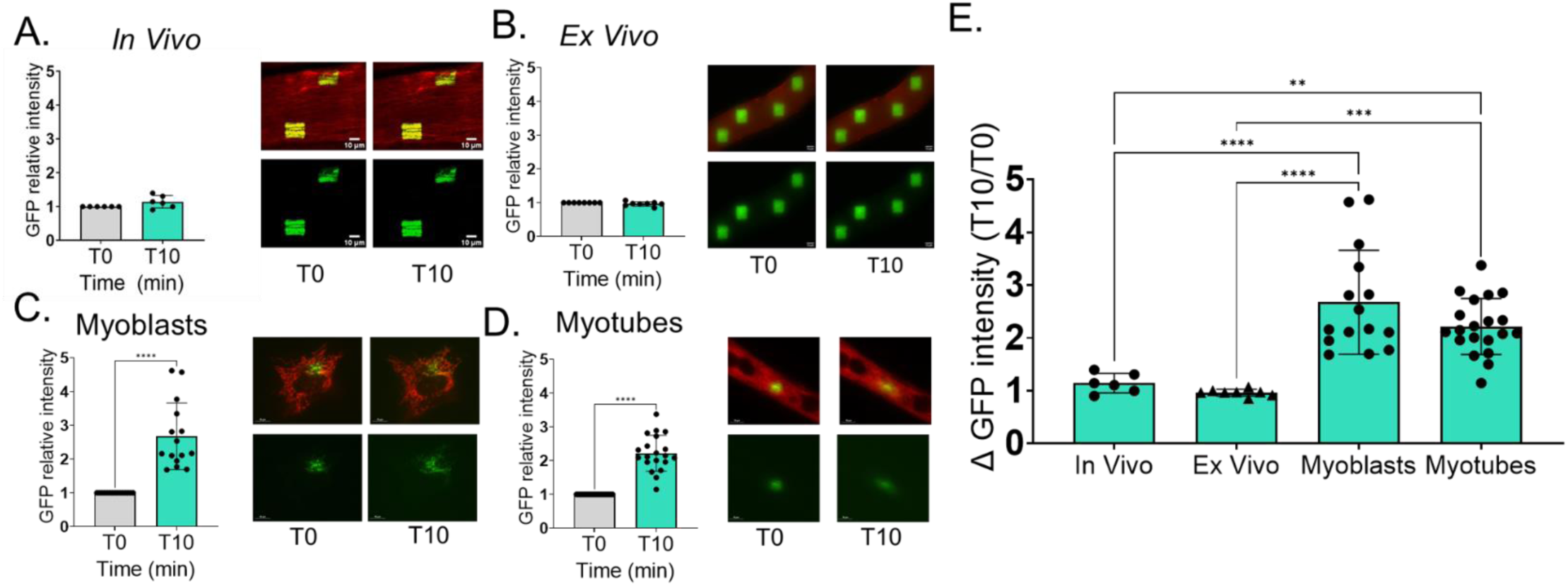
*In vivo photoactivation* imaging compared to *ex vivo* and *in vitro models.* The photoactivatable GFP intensity spread at T0 immediately after photoactivation (gray bar) and at T10 GFP spread 10 minutes post photoactivation normalized to T0 (green bar) and representative images of mitochondrial targeted mScarlet (red) and photoactivatable GFP (green) of **(A)** *In Vivo* Tibialis Anterior of young mice (n=6). Each image consisted of a 35-step z-stack (1024×1024 pixels) and were acquired through a 25 x 1.1 NA Nikon water objective with 3x zoom using Nikon NIS elements. ROIs were activated using the Coherent Chameleon Vision Ultra tunable IR laser tuned to 760 nm, with 10 loops at 1% laser power (Scale= 10 µM) **(B)** *Ex Vivo* single fibers isolated from the TA of young mice (n=8). Each image consisted of a 35-step z-stack (2304×2304 pixels) and were acquired through a 40x on a Nikon Ti2-E microscope. (Scale= 14 µM) ROIs were drawn and assigned to a stimulation protocol carried out with an Optimicroscan XY galvo scanning unit (405nm stimulation laser at 2% laser power for 100µs per pixel). **(C)** *In Vitro* Myoblasts and **(D)** *In Vitro* Myotubes. Each image consisted of a single plane image (2304×2304 pixels) and were acquired through a 60x oil immersion objective in a stage top incubator using a Nikon Ti2-E microscope using the same ROI activation as B (Scale= 10 µM). **(E)** Comparison of four models of GFP spread normalized to T0 spread. Contrast of all images were increased for visual clarity only. Asterisks indicate a statistical difference. T-tests were used to compare T0 and T10 and a One-Way ANOVA was completed for comparison of the four models, p<0.5. Data are presented as means ± SD.

**Figure 4.**
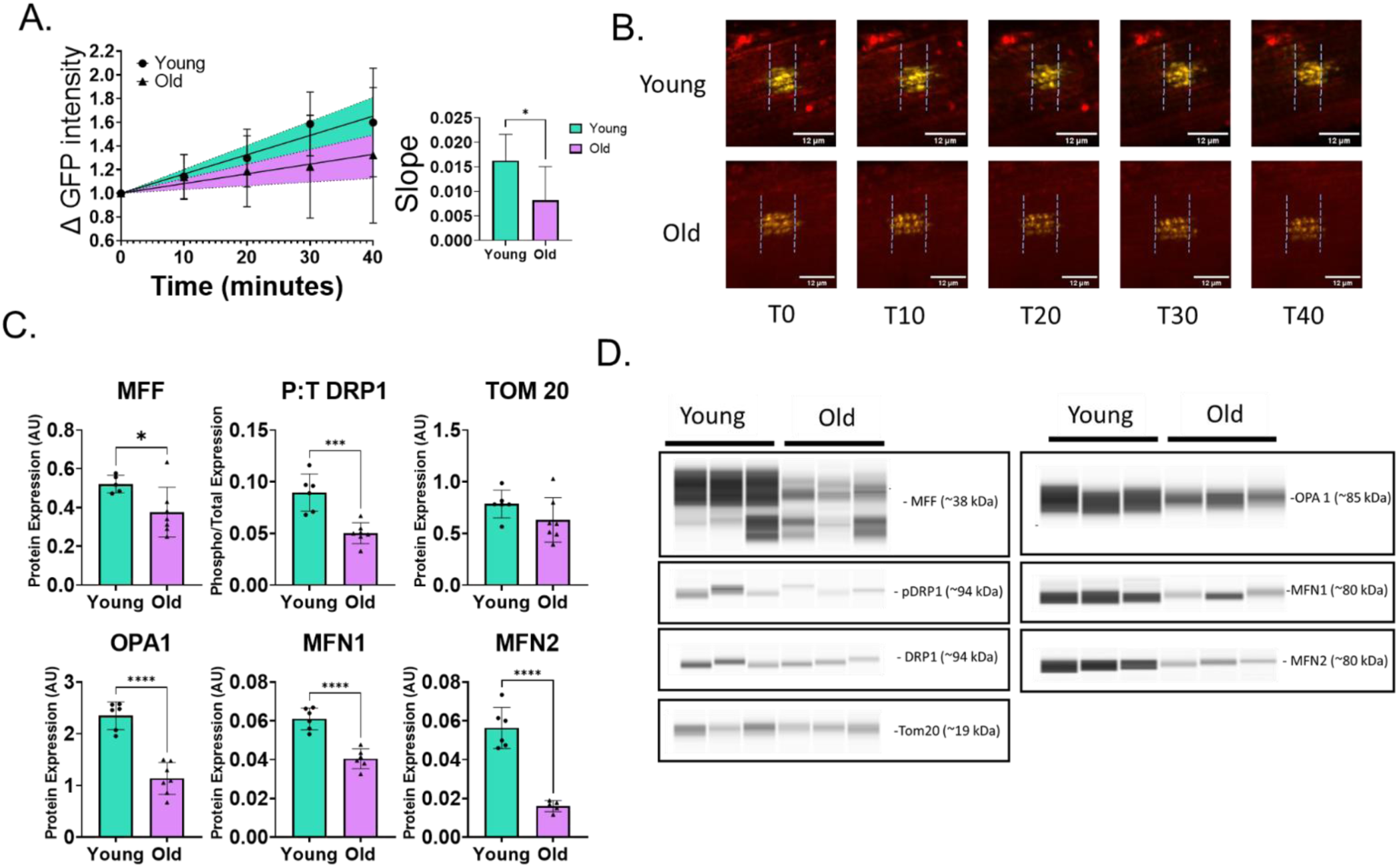
*In vivo photoactivation* imaging of old and young mice. **(A)** Change in GFP intensity spread over 40 min comparing young (circle) and old (triangle) Tibialis Anterior with 95% confidence intervals of young (green) and old (purple). Slope of the line for the change in GFP intensity spread over 40 min for young (green) and old (purple). **(B)** Representative images of GFP intensity spread (green) for young and old animals from T0 to T40 (scale = 12µM). Dotted lines indicate edge of photoactivated area at T0. Each image consisted of a 35-step z-stack (1024×1024 pixels) and were acquired through a 25x 1.1 NA Nikon water objective with 3x zoom using Nikon NIS elements (Nikon instruments Inc., NY, USA. Brightness and contrast adjusted equally across all images for visual clarity only. **(C)** Comparison of young and old TA protein expression of fission and fusion markers normalized to total protein in arbitrary units. **(D)** Representative images of the protein simple capillary blots with molecular weights. T-tests were completed to compare young and old animals. Data are presented as means ± SD. Asterisks indicate p<0.05.

Next, we used the same imaging technique with a longer period of imaging to compare the rate of mitochondrial dynamic events of young (6 months) versus old (24 months) mice. There was a main effect of time with the spread of GFP increasing over time in the young group **(Figures 4a and 4b).** Over time there was increased variability in GFP spread, but overall, there were no differences in the GFP spread from T0 to T40 in the old animals. A comparison of the slope of the two lines demonstrates the lack of GFP spread in the old animals **(Figure 4a).** The slope of GFP spread in the young animals was two times higher (0.016 ± 0.002 events/min) than the old animals (0.008 ± 0.003 events/min), indicating faster rates of dynamic events.

Markers of mitochondrial fission and fusion are consistently used to assess mitochondrial dynamics despite their static nature. To determine whether markers of fusion and fission reflected measured rates of mitochondrial dynamics *in vivo,* we compared the *in vivo* imaging data to markers of mitochondrial dynamics. We selected several proteins involved in fission (DRP1, MFF) and fusion (OPA1, MFN1, MFN2) **(Figure 4C and 4D).** Overall, the TA of old animals had lower expression of all fission and fusion proteins assayed. For fusion proteins, old animals had two times lower expression of OPA1 than the young animals. Old animals also had lower expression of MFN1 and MFN2 than the young animals. Old animals had a lower expression of MFF and the protein expression of the phosphorylated (Ser616) to total ratio of DRP1 was also lower in the old animals. However, there was no difference between the old and young animals for TOM20, which is often used as a measure of mitochondrial content, volume, or mass (42–44). This finding suggests a reduction in the protein abundance of fission and fusion proteins in old mice rather than an overall reduction in total mitochondrial content.

Previous literature indicates that both acute and chronic exercise can stimulate mitochondrial dynamics (16, 17, 45). To assess whether acute exercise can impact the rate of mitochondrial dynamics in young and old animals as predicted, we measured TA mitochondrial dynamics *in vivo* after a single treadmill bout. While the young exercise animals showed the same trend as their sedentary counterparts of increased GFP spread over time, there was no difference between the young sedentary and exercised animals (**Figure 5A and 5B)** indicating that the acute bout of exercise did not increase the rate of dynamic events in young animals. However, there was a main effect of time with GFP spread increasing over time in the old exercise animals, suggesting that the rate of mitochondrial dynamics increased after the acute bout of exercise. The old exercise group had a higher slope than their old counterparts but was not different from the young animal groups. This higher rate of GFP spread in the old exercise group indicates that acute exercise increased mitochondrial dynamic rates in the old animals.

**Figure 5.**
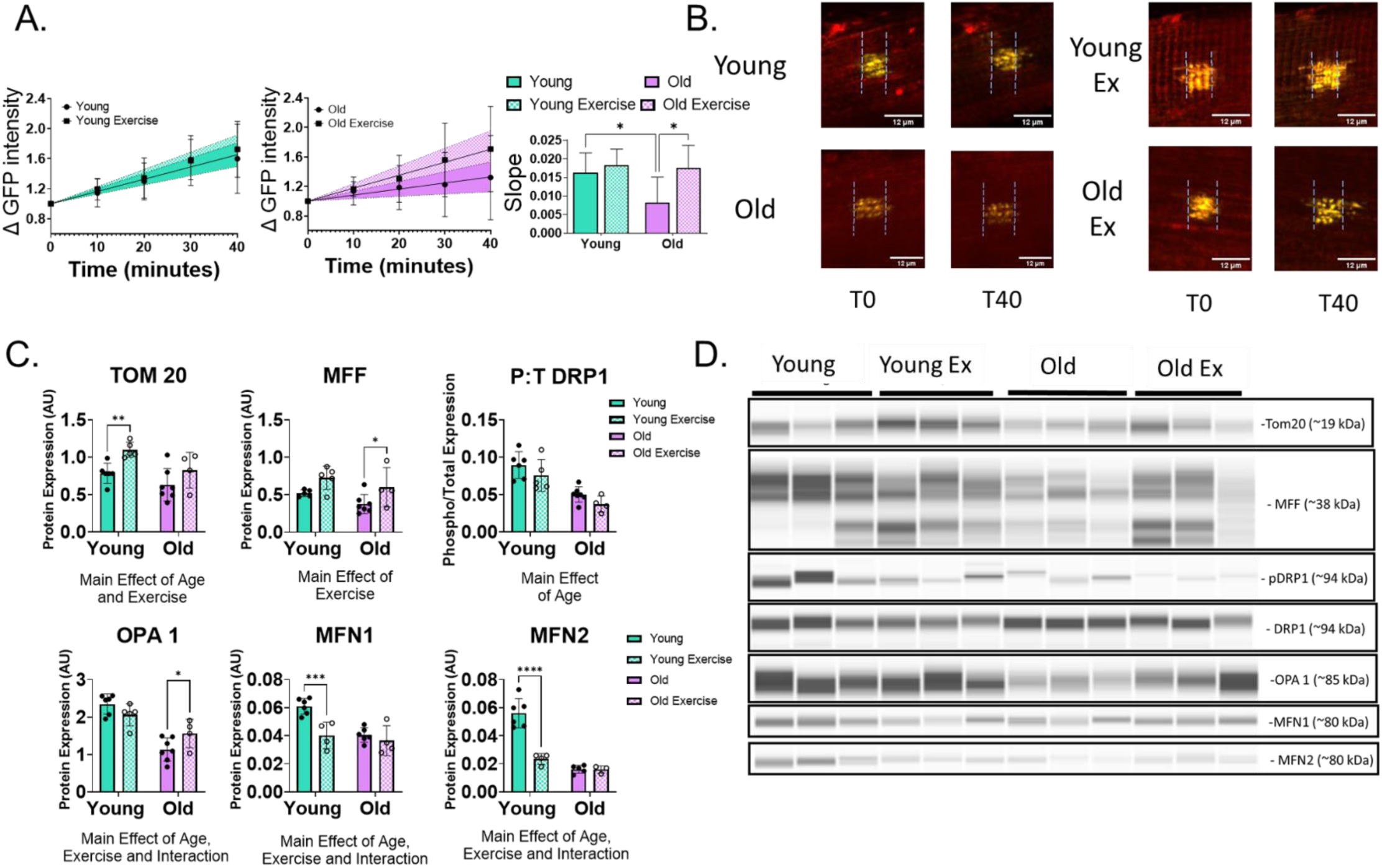
*In vivo photoactivation* imaging of old and young exercised mice. **(A)** Change in GFP intensity spread over 40 min of young (circle), young exercised (square), old (circle) and old exercise (square) Tibialis Anterior with 95% confidence intervals of young (dark green), young exercise (lighter green), old (dark purple) and old exercise (light purple). Slope of the lines of the change in GFP intensity spread over 40 min for young (green), young exercise (light green), old (purple) and old exercise (light purple) **(B)** Representative images of GFP intensity spread (green) for young, young exercise, old and old exercise at T0 and T40 (scale = 12µM). Dotted lines indicate edge of photoactivated area at T0. Each image consisted of a 35-step z-stack (1024×1024 pixels) and were acquired through a 25x 1.1 NA Nikon water objective with 3x zoom using Nikon NIS elements (Nikon instruments Inc., NY, USA) Brightness and contrast adjusted equally across all images for visual clarity only. **(C)** Comparison of young, young exercise, old and old exercise TA protein expression of fission and fusion markers normalized to total protein in arbitrary units. **(D)** Representative images of the protein simple capillary blots. One-way ANOVA were performed to compare the four groups. Data are presented as means ± SD. Asterisks indicate p<0.05.

We compared our *in vivo* responses to exercise to markers of fusion and fission (**Figures 5C and 5D)** to determine if protein markers reflect the measured dynamic rates after an acute bout of exercise. Protein expression analysis showed a main effect of age and exercise on TOM20, with an increase of TOM20 protein in the young exercised animals and a similar trend (p=0.09) in the old animals. TOM20 is commonly used as a marker of mitochondrial content; however, these findings in isolation could misleadingly suggest changes in overall mitochondrial content following an acute bout of exercise. A main effect of age and exercise was found in all three of the fusion markers. Muscles of old exercised animals expressed more OPA1 protein than their sedentary counterparts, but overall, the old animals had lower OPA1 protein than the young (p<0.05). Young exercised animals had less MFN1 and MFN2 protein than young sedentary animals (p<0.05), but there were no differences between exercise and sedentary old animals. With regards to fission proteins, the exercised old group was greater than sedentary old group for MFF and resulted in a similar trend in young animals (p=0.052). The phosphorylation state of DRP1 was lower in old animals but with no differences between the sedentary and exercised animals at either age. Overall, protein analysis provided no clear consensus regarding a change in mitochondrial dynamics following an acute exercise bout.

## Discussion

We developed a 2-photon imaging approach coupled with matrix-targeted fluorophores to measure *in vivo* rates of mitochondrial dynamics remodeling events in skeletal muscle. Our investigation showed that the rate of dynamic events *in vivo* is slower than previously indicated by *in vitro* models. However, our *ex vivo* analyses demonstrated similar rates to our *in vivo* rates. These findings underscore the disparities between murine cell lines versus animal models suggesting that extrapolating *in vitro* findings to *in vivo* conditions may overestimate the rates of fusion and fission events. Moreover, we validated that our method is sensitive enough to measure differences between experimental conditions such as age and exercise despite slower rates of dynamics than initially expected. We observed that dynamic rates are slower in old sedentary animals than in young sedentary animals. Furthermore, we observed a higher rate of dynamics in old exercised animals compared to their sedentary counterparts, while young exercised animals showed no difference from their sedentary controls. Finally, our data provide compelling evidence that drawing conclusions solely from static markers of mitochondrial dynamics may be misleading and warrants careful consideration.

Despite the understanding that fusion and fission are dynamic processes that continually respond to the energetic needs of the tissue, current methods to assess mitochondrial dynamics primarily rely on single time point (static) measures. The most common and arguably the most accessible of these static measures is the use of protein expression level at a particular time point as a correlate for activity or markers involved in either fusion or fission (18, 42, 46, 47). Similarly, researchers employ electron microscopy or other visual mitochondrial morphology measures at a distinct time point to assess mitochondrial shape and provide information on the balance between fusion and fission at a particular time experimentally (48–51). While several groups have employed multi-time point assessment in cultured models of skeletal muscle (26, 52), to our knowledge only one study (32) has attempted *in vivo* imaging in skeletal muscle.

Our 2-photon imaging approach built upon the studies mentioned above (26, 32, 52) and allowed us to measure real-time changes in dynamic events. Like many other cellular processes, we hypothesized that the rate of mitochondrial dynamics events, would be slower *in vivo* than *ex vivo* and *in vitro*. This hypothesis was mainly driven by significant differences in environment and our previous findings that *in vitro* protein turnover, another essential feature of mitochondrial remodeling, is higher in C2C12 (29) than *in vivo* models (21, 41, 53). We found similar rates of dynamics in our single fibers *ex vivo* when compared to our *in vivo* samples **(Figure 2)**. Our findings agree with previous data in cultured muscle fibers from the EDL (27), which used a confocal imaging process. This study found that fusion events were sparse in the more glycolytic type 2b single fibers over a 30-minute time period (27). The EDL and TA of the mouse are both primarily glycolytic muscles (54), and Mishra et al. (2015) found that the more glycolytic fibers had isolated, punctate mitochondria, while the oxidative fiber types had more elongated and interconnected mitochondria with higher rates of fusion (27). Our *in vivo/ex vivo* comparison indicates that under some conditions that do not require repeated measurements, *ex vivo* approaches could be used as a surrogate for *in vivo* analysis.

In contrast to our *ex vivo* findings, myoblasts and myotubes had much faster rates of mitochondrial dynamic events than our *in vivo* model **(Figure 3).** We do not consider myoblasts a representative model of mature skeletal muscle because they lack the underlying structure of skeletal muscle fibers. However, C2C12 myotubes are often used as a surrogate model to understand *in vivo* outcomes. It was important to establish if the rates of mitochondrial dynamics in myotubes reflected what is occurring *in vivo*. While we did not specifically measure individual events, the rate of the spread of photo-activated fluorescent signal over time agreed with previous work in primary cultures of myofibers that found 0.5 fusion events and 0.15 fission events per minute (26). In our *in vivo* model we found rates ranging from 0.08 dynamic events per minute to 0.16 dynamic events per minute. The ability to compare our *in vitro* results to our *in vivo* model provides a better understanding of the differences between experimental models and the conclusions we can make about dynamic rates in these different models.

The critical role that mitochondrial dynamics play in all aspects of mitochondrial function makes them important targets of aging skeletal muscle research. While numerous studies have explored the impact of aging on the regulation of mitochondrial dynamics (as reviewed in 20), a consensus regarding the balance or regulation of these forces remains elusive. This lack of conclusive evidence, driven mainly by the reliance on static markers, suggests that an approach relying on snapshots of a dynamic process may not tell us the whole story. Various factors such as circadian rhythm, feeding, and time since last bout of physical activity could all influence the rates of dynamic events. We have observed that rates of mitochondrial dynamics in an unperturbed skeletal muscle are relatively slow, but the skeletal muscle may undergo transient periods of rapid remodeling depending on the conditions it experiences. Without longitudinal assessments, there is little ability to discern if and when these factors impact mitochondrial dynamics, potentially leading to misguided or erroneous conclusions. Therefore, we must incorporate longitudinal and direct measures of mitochondrial dynamics to gain deeper insights into the physiology of these organelles.

When comparing young to old animals **(Figure 4),** our protein marker data suggests that both fusion and fission are lower in the old animals, a trend generally consistent with our *in vivo* imaging results. However, considering the protein marker data without real-time imaging could lead to different conclusions based on the magnitudes of differences between young and old protein expression. Several studies have suggested that aging skeletal muscle mitochondria may preferentially undergo fission, with elevated levels of FIS1 and DRP1 and downregulated MFN2 (49, 54). More recently, others have found morphology indicative of higher fusion levels with no significant differences in fusion and fission protein markers (56). These inconsistencies could be explained by various aspects of experimental design including muscle type, experimental model, and timing of assessments. Additionally, skeletal muscle contains distinct pools of mitochondria, subsarcolemmal and intermyofibrillar, which are challenging to differentiate without spatial information within the muscle fibers, yet have functional differences (57). Our new *in vivo* technique would allow for the comparison of dynamic rates between these distinct mitochondrial pools, offering a distinct advantage over *ex vivo* and static measures. Given the inconsistencies in previous literature and the complexities of mitochondria in skeletal muscle, caution is warranted when drawing conclusions based solely on static assessments of a clearly dynamic process.

The response to an exercise bout further illustrated a need for caution when using protein markers **(Figure 5)**.In our study there was minimal consensus among the markers on the influence of an acute bout of exercise on fusion and fission. Previous findings have suggested that physically active subjects have augmented fusion with long-term exercise in whole muscle extracts (17, 18, 44). Additionally, it has been suggested that acute exercise can enhance MFN2 mRNA expression in young subjects (16) and that this enhanced fusion could assist in mitochondrial function of older individuals. Despite the generalized idea that aerobic exercise enhances fusion, a recent study found that both markers of fusion and fission were increased in active young and old male subjects (57), indicating that fusion and fission acted in complementary ways to enhance mitochondrial function. Disparities across studies may again arise from various experimental design aspects. Yet, when contrasted with our *in vivo* imaging data, they highlight how markers alone can be misleading, causing potential confusion and contradictions.

There are limitations to the current study. The first is that our analysis does not differentiate between fusion and fission events. Our primary interest was developing a method that could make measurements over a relatively long period of time *in vivo* that could assess the overall rate of mitochondrial dynamics, which we have accomplished. While we cannot differentiate the fission or fusion events, our data and data from others (26) strongly suggest that fusion dominates during remodeling events in densely packed networks like skeletal muscle. Second, the sample sizes of our exercise groups were small, yet we did see differences between the old and old exercised animals. Since there were no previous studies to base *a priori* power calculations on, this study establishes expected variability, both during sedentary conditions and with an intervention, for future studies. Next, our hypothesis that rates of dynamic events would be slower *in vivo* compared to *in vitro* models was correct, although our measured *in vivo* rates were slower than we expected. Keeping animals under anesthesia for long periods impacts many aspects of physiology and is especially hard on old animals (59, 60). Due to the slow rates of dynamic events *in vivo*, measuring the number of events and distinguishing between fusion and fission events may require more time under anesthesia than would be desirable.

In summary, our novel method using multiphoton microscopy demonstrates it is possible to image mitochondrial dynamics *in vivo.* Our *in vivo* method requires specialized equipment that may not be available at all institutions, but the technique itself is simple and easy to implement. The imaging process is time consuming due to the slow rates of mitochondrial dynamics in skeletal muscle but appropriately captures rates of remodeling. Through imaging both mitochondrial calcium sensors and mitochondrial-targeted photoactivatable GFP tags, we have validated that this technique does not influence mitochondrial function and is sensitive enough to distinguish changes of various magnitudes and speeds. As expected, the dynamic rates were slower *in vivo* compared to *in vitro* models. However, the unexpected finding that single fibers isolated from animals were similar to the *in vivo* model provides a reasonably logical alternative for longitudinal assessment when *in vivo* imaging is not possible. Additionally, our study found that markers of mitochondrial dynamics do not always reflect dynamic rates *in vivo*, and interpretation of changes in mitochondrial dynamics using static measures alone has limitations. We chose to validate this method in the TA, but this technique is not limited to the TA and has potential application in other superficial muscles like the gastrocnemius. While there are many aspects of this technique to develop and improve upon, the ability to provide longitudinal information about the critical functions of fusion and fission in skeletal muscle will help answer many experimental questions about the role of mitochondrial dynamics in skeletal muscle aging and disease.

## Data availability

All Source data is provided on a figshare repository https://doi.org/10.6084/m9.figshare.c.7482933.v1. Raw versions of all images presented in figures are also provided. All other images are available upon reasonable request.

## Author Contributions

CLO, AD, BFM and TLL were responsible for conception and design of the experiments. CLO, AD, KC, SD, MB, and AB acquired and analyzed data for this work. CLO, TLL, and BFM drafted the work, and all authors critically revised this paper.

## Funding

NIH T32AG052363 – CLO

NIH R35GM137921 – TLL

Presbyterian Health Foundation – TLL & BFM

## Acknowledgements

We thank members of the Lewis and Miller labs for feedback and helpful discussion.

---

**Supplemental Figure 1:**
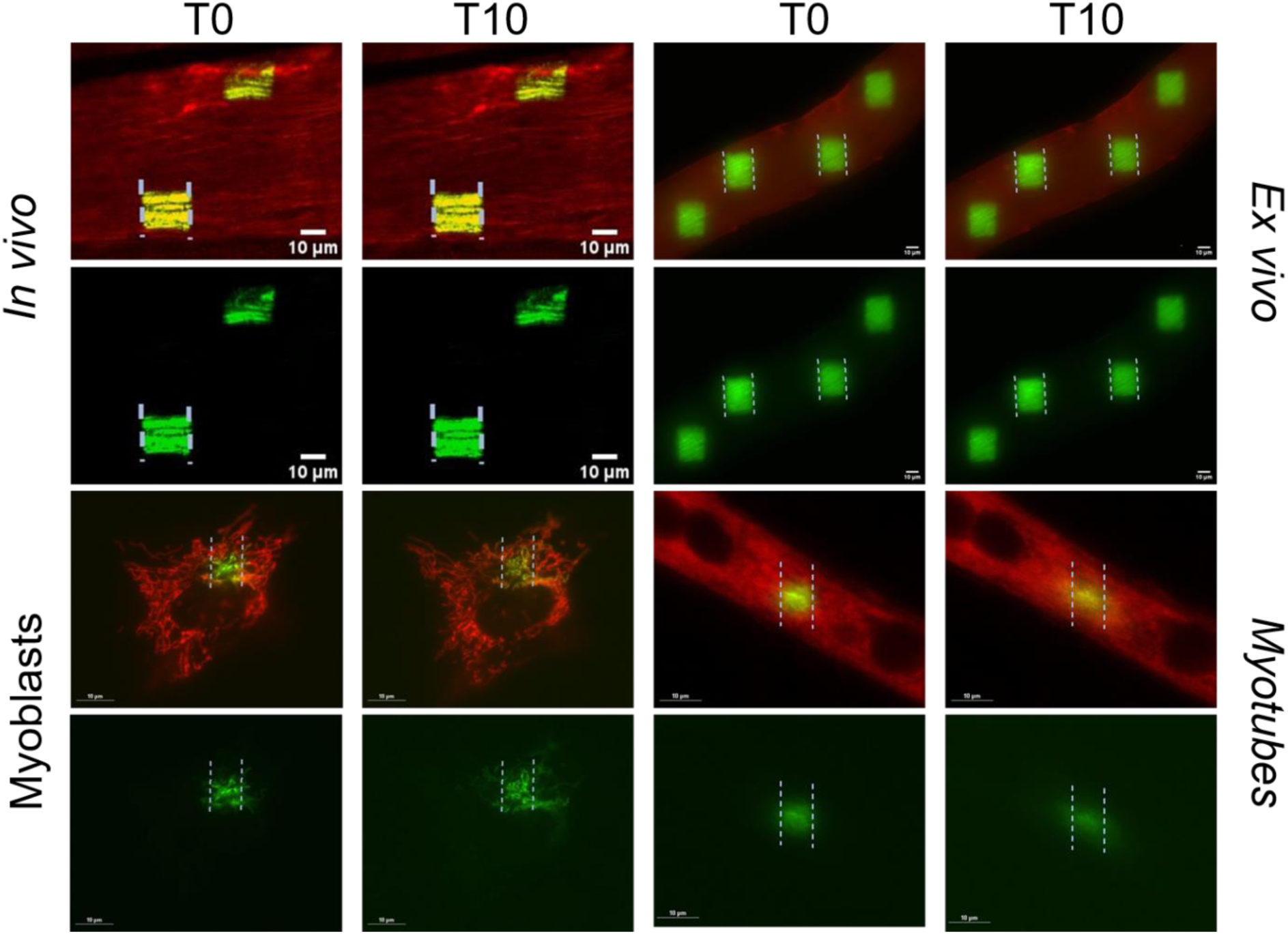
Enlarged photos of *In vivo photoactivation* imaging compared to *ex vivo* and *in vitro models.* Representative images of the photoactivatable GFP intensity spread immediately after photoactivation (T0) and GFP spread 10 minutes post photoactivation (T10) using electroporated fluorophores mitochondrial targeted mScarlet (red) and photoactivatable GFP (green) of *in vivo* tibialis anterior of young mice (n=6). Each image consisted of a 35-step z-stack (1024×1024 pixels) and were acquired through a 25 x 1.1 NA Nikon water objective with 3x zoom using Nikon NIS elements. ROIs were activated using the Coherent Chameleon Vision Ultra tunable IR laser tuned to 760 nm, with 10 loops at 1% laser power (Scale= 10 µM). *Ex Vivo* single fibers isolated from the TA of young mice (n=8). Each image consisted of a 35-step z-stack (2304×2304 pixels) and were acquired through a 40x on a Nikon Ti2-E microscope. (Scale= 14 µM) ROIs were drawn and assigned to a stimulation protocol carried out with an Optimicroscan XY galvo scanning unit (405nm stimulation laser at 2% laser power for 100µs per pixel). *In Vitro* Myoblasts and *In Vitro* Myotubes. Each image consisted of a single plane image (2304×2304 pixels) and were acquired through a 60x oil immersion objective in a stage top incubator using a Nikon Ti2-E microscope using the same ROI activation as B (Scale= 10 µM). Contrast of all images were increased for visual clarity only.

**Supplemental Figure 2:**
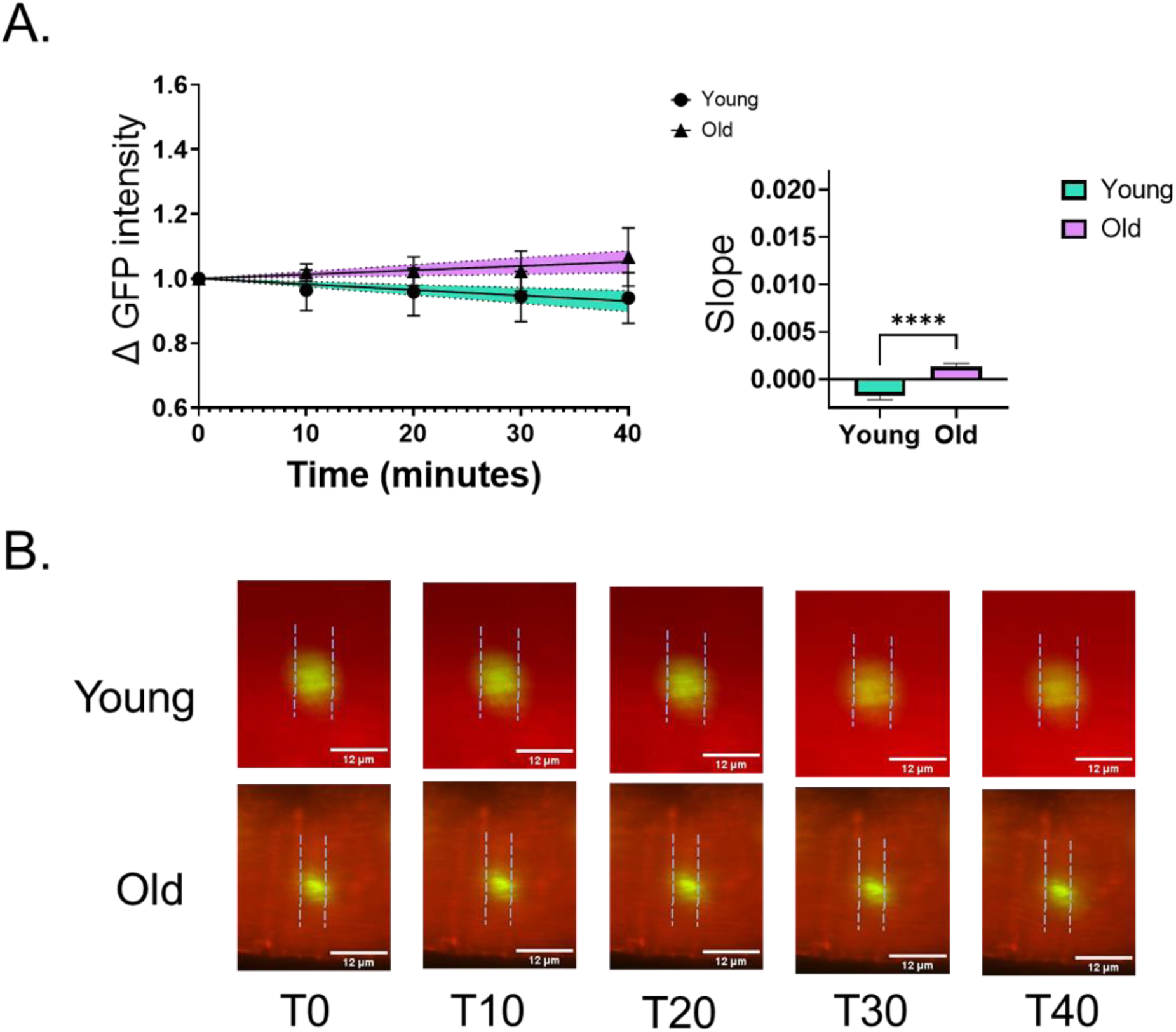
*Ex vivo photoactivation* imaging of old and young isolated fibers. **(A)** Change in GFP intensity spread over 40 min comparing young (circle) and old (triangle) isolated tibialis anterior fibers with 95% confidence intervals of young (green) and old (purple). Slope of the line for the change in GFP intensity spread over 40 min for young (green) and old (purple). **(B)** Representative images of change in GFP intensity spread (green) for young and old animals from T0 to T40 (scale = 12µM). Dotted lines indicate edge of photoactivated area at T0. Each image consisted of a 35-step z-stack (2304×2304 pixels) and were acquired through a 40x Nikon on a Ti2-E microscope. ROIs were drawn and assigned to a stimulation protocol carried out with an Optimicroscan XY galvo scanning unit (405nm stimulation laser at 2% laser power for 100µs per pixel). Brightness and contrast adjusted equally across all images for visual clarity only. Data are presented as means ± SD.

## References

1. dos Santos, L., Cyrino, E. S., Antunes, M., Santos, D. A., and Sardinha, L. B. (2017) Sarcopenia and physical independence in older adults: the independent and synergic role of muscle mass and muscle function. Journal of Cachexia, Sarcopenia and Muscle 8, 245– 250

2. Xu, W., Chen, T., Cai, Y., Hu, Y., Fan, L., and Wu, C. (2020) Sarcopenia in Community-Dwelling Oldest Old is Associated with Disability and Poor Physical Function. J Nutr Health Aging 24, 339–345

3. Janssen, I., Shepard, D. S., Katzmarzyk, P. T., and Roubenoff, R. (2004) The Healthcare Costs of Sarcopenia in the United States. Journal of the American Geriatrics Society 52, 80–85

4. Larsson, L., Degens, H., Li, M., Salviati, L., Lee, Y. I., Thompson, W., Kirkland, J. L., and Sandri, M. (2019) Sarcopenia: Aging-Related Loss of Muscle Mass and Function. Physiological Reviews 99, 427–511

5. Short, K. R., Bigelow, M. L., Kahl, J., Singh, R., Coenen-Schimke, J., Raghavakaimal, S., and Nair, K. S. (2005) Decline in skeletal muscle mitochondrial function with aging in humans. Proceedings of the National Academy of Sciences 102, 5618–5623

6. Musci, R. V., Hamilton, K. L., and Miller, B. F. (2018) Targeting mitochondrial function and proteostasis to mitigate dynapenia. Eur J Appl Physiol 118, 1–9

7. Coen, P. M., Musci, R. V., Hinkley, J. M., and Miller, B. F. (2019) Mitochondria as a Target for Mitigating Sarcopenia. Frontiers in Physiology 9

8. Kirkwood, S. P., Munn, E. A., and Brooks, G. A. (1986) Mitochondrial reticulum in limb skeletal muscle. Am J Physiol 251, C395–402

9. Kirkwood, S. P., Munn, E. A., and Brooks, G. A. (1986) Mitochondrial reticulum in limb skeletal muscle. American Journal of Physiology-Cell Physiology 251, C395–C402

10. Hoppeler, H., Hudlicka, O., and Uhlmann, E. (1987) Relationship between mitochondria and oxygen consumption in isolated cat muscles. J Physiol 385, 661–675

11. Ogata, T. and Yamasaki, Y. (1997) Ultra-high-resolution scanning electron microscopy of mitochondria and sarcoplasmic reticulum arrangement in human red, white, and intermediate muscle fibers. Anat Rec 248, 214–223

12. Glancy, B., Hartnell, L. M., Malide, D., Yu, Z.-X., Combs, C. A., Connelly, P. S., Subramaniam, S., and Balaban, R. S. (2015) Mitochondrial reticulum for cellular energy distribution in muscle. Nature 523, 617–620

13. Glancy, B., Kim, Y., Katti, P., and Willingham, T. B. (2020) The Functional Impact of Mitochondrial Structure Across Subcellular Scales. Frontiers in Physiology 11

14. Lee, Y., Jeong, S.-Y., Karbowski, M., Smith, C. L., and Youle, R. J. (2004) Roles of the mammalian mitochondrial fission and fusion mediators Fis1, Drp1, and Opa1 in apoptosis. Mol Biol Cell 15, 5001–5011

15. Chen, H., Detmer, S. A., Ewald, A. J., Griffin, E. E., Fraser, S. E., and Chan, D. C. (2003) Mitofusins Mfn1 and Mfn2 coordinately regulate mitochondrial fusion and are essential for embryonic development. J Cell Biol 160, 189–200

16. Cartoni, R., Léger, B., Hock, M. B., Praz, M., Crettenand, A., Pich, S., Ziltener, J.-L., Luthi, F., Dériaz, O., Zorzano, A., Gobelet, C., Kralli, A., and Russell, A. P. (2005) Mitofusins 1/2 and ERRα expression are increased in human skeletal muscle after physical exercise. The Journal of Physiology 567, 349–358

17. Konopka, A. R., Suer, M. K., Wolff, C. A., and Harber, M. P. (2014) Markers of Human Skeletal Muscle Mitochondrial Biogenesis and Quality Control: Effects of Age and Aerobic Exercise Training. The Journals of Gerontology: Series A 69, 371–378

18. Arribat, Y., Broskey, N. T., Greggio, C., Boutant, M., Conde Alonso, S., Kulkarni, S. S., Lagarrigue, S., Carnero, E. A., Besson, C., Cantó, C., and Amati, F. (2019) Distinct patterns of skeletal muscle mitochondria fusion, fission and mitophagy upon duration of exercise training. Acta Physiologica 225, e13179

19. Mao, X., Gu, Y., Sui, X., Shen, L., Han, J., Wang, H., Xi, Q., Zhuang, Q., Meng, Q., and Wu, G. (2021) Phosphorylation of Dynamin-Related Protein 1 (DRP1) Regulates Mitochondrial Dynamics and Skeletal Muscle Wasting in Cancer Cachexia. Front Cell Dev Biol 9, 673618

20. Leduc-Gaudet, J.-P., Hussain, S. N. A., Barreiro, E., and Gouspillou, G. (2021) Mitochondrial Dynamics and Mitophagy in Skeletal Muscle Health and Aging. International Journal of Molecular Sciences 22, 8179

21. Miller, B. F., Robinson, M. M., Bruss, M. D., Hellerstein, M., and Hamilton, K. L. (2012) A comprehensive assessment of mitochondrial protein synthesis and cellular proliferation with age and caloric restriction. Aging Cell 11, 150–161

22. Lewis, T. L., Turi, G. F., Kwon, S. K., Losonczy, A., and Polleux, F. (2016) Progressive Decrease of Mitochondrial Motility during Maturation of Cortical Axons In Vitro and In Vivo. Current Biology 26, 2602–2608

23. Magrané, J., Cortez, C., Gan, W.-B., and Manfredi, G. (2014) Abnormal mitochondrial transport and morphology are common pathological denominators in SOD1 and TDP43 ALS mouse models. Hum Mol Genet 23, 1413–1424

24. Pouli, D., Balu, M., Alonzo, C. A., Liu, Z., Quinn, K. P., Rius-Diaz, F., Harris, R. M., Kelly, K. M., Tromberg, B. J., and Georgakoudi, I. (2016) Imaging mitochondrial dynamics in human skin reveals depth-dependent hypoxia and malignant potential for diagnosis. Science Translational Medicine 8

25. Lewis, T. L., Kwon, S. K., Lee, A., Shaw, R., and Polleux, F. (2018) MFF-dependent mitochondrial fission regulates presynaptic release and axon branching by limiting axonal mitochondria size. Nature Communications 9, 1–15

26. Eisner, V., Lenaers, G., and Hajnóczky, G. (2014) Mitochondrial fusion is frequent in skeletal muscle and supports excitation-contraction coupling. Journal of Cell Biology 205, 179–195

27. Mishra, P., Varuzhanyan, G., Pham, A. H., and Chan, D. C. (2015) Mitochondrial Dynamics Is a Distinguishing Feature of Skeletal Muscle Fiber Types and Regulates Organellar Compartmentalization. Cell Metabolism 22, 1033–1044

28. Dörrbaum, A. R., Kochen, L., Langer, J. D., and Schuman, E. M. (2018) Local and global influences on protein turnover in neurons and glia. eLife 7, e34202

29. Wolff, C. A., Reid, J. J., Musci, R. V., Bruns, D. R., Linden, M. A., Konopka, A. R., Peelor, F. F., Miller, B. F., and Hamilton, K. L. (2020) Differential Effects of Rapamycin and Metformin in Combination With Rapamycin on Mechanisms of Proteostasis in Cultured Skeletal Myotubes. J Gerontol A Biol Sci Med Sci 75, 32–39

30. Price, J. C., Guan, S., Burlingame, A., Prusiner, S. B., and Ghaemmaghami, S. (2010) Analysis of proteome dynamics in the mouse brain. Proc Natl Acad Sci U S A 107, 14508– 14513

31. Romanello, V., Guadagnin, E., Gomes, L., Roder, I., Sandri, C., Petersen, Y., Milan, G., Masiero, E., Del Piccolo, P., Foretz, M., Scorrano, L., Rudolf, R., and Sandri, M. (2010) Mitochondrial fission and remodelling contributes to muscle atrophy. EMBO J 29, 1774– 1785

32. Liu, R., Jin, P., LiqunYu, Wang, Y., Han, L., Shi, T., and Li, X. (2014) Impaired Mitochondrial Dynamics and Bioenergetics in Diabetic Skeletal Muscle. PLOS ONE 9, e92810

33. Benninger, R. K. P. and Piston, D. W. (2013) Two-Photon Excitation Microscopy for the Study of Living Cells and Tissues. Curr Protoc Cell Biol 0 4, Unit-4.1124

34. Hughes, D. C., Hardee, J. P., Waddell, D. S., and Goodman, C. A. (2022) CORP: Gene delivery into murine skeletal muscle using in vivo electroporation. Journal of Applied Physiology 133, 41–59

35. Keire, P., Shearer, A., Shefer, G., and Yablonka-Reuveni, Z. (2013) Isolation and Culture of Skeletal Muscle Myofibers as a Means to Analyze Satellite Cells. In Basic Cell Culture Protocols (Helgason, C. D. and Miller, C. L., eds) pp. 431–468, Humana Press, Totowa, NJ

36. He, W. A., Calore, F., Londhe, P., Canella, A., Guttridge, D. C., and Croce, C. M. (2014) Microvesicles containing miRNAs promote muscle cell death in cancer cachexia via TLR7. Proceedings of the National Academy of Sciences 111, 4525–4529

37. Velayuthan, L. P., Moretto, L., Tågerud, S., Ušaj, M., and Månsson, A. (2023) Virus-free transfection, transient expression, and purification of human cardiac myosin in mammalian muscle cells for biochemical and biophysical assays. Sci Rep 13, 4101

38. Miller, B., Hamilton, K., Boushel, R., Williamson, K., Laner, V., Gnaiger, E., and Davis, M. (2017) Mitochondrial respiration in highly aerobic canines in the non-raced state and after a 1600-km sled dog race. PLOS ONE 12, e0174874

39. Konopka, A. R., Castor, W. M., Wolff, C. A., Musci, R. V., Reid, J. J., Laurin, J. L., Valenti, Z. J., Hamilton, K. L., and Miller, B. F. (2017) Skeletal muscle mitochondrial protein synthesis and respiration in response to the energetic stress of an ultra-endurance race. J Appl Physiol (1985) 123, 1516–1524

40. Bubak, M. P., Davidyan, A., O’Reilly, C. L., Mondal, S. A., Keast, J., Doidge, S. M., Borowik, A. K., Taylor, M. E., Volovičeva, E., Kinter, M. T., Britton, S. L., Koch, L. G., Stout, M. B., Lewis Jr, T. L., and Miller, B. F. (2024) Metformin treatment results in distinctive skeletal muscle mitochondrial remodeling in rats with different intrinsic aerobic capacities. Aging Cell 23, e14235

41. Miller, B. F., Wolff, C. A., Peelor, F. F., Shipman, P. D., and Hamilton, K. L. (2015) Modeling the contribution of individual proteins to mixed skeletal muscle protein synthetic rates over increasing periods of label incorporation. J Appl Physiol (1985) 118, 655–661

42. Buso, A., Comelli, M., Picco, R., Isola, M., Magnesa, B., Pišot, R., Rittweger, J., Salvadego, D., Šimunič, B., Grassi, B., and Mavelli, I. (2019) Mitochondrial Adaptations in Elderly and Young Men Skeletal Muscle Following 2 Weeks of Bed Rest and Rehabilitation. Front. Physiol. 10

43. Nijholt, K. T., Meems, L. M. G., Ruifrok, W. P. T., Maass, A. H., Yurista, S. R., Pavez-Giani, M. G., Mahmoud, B., Wolters, A. H. G., van Veldhuisen, D. J., van Gilst, W. H., Silljé, H. H. W., de Boer, R. A., and Westenbrink, B. D. (2021) The erythropoietin receptor expressed in skeletal muscle is essential for mitochondrial biogenesis and physiological exercise. Pflugers Arch - Eur J Physiol 473, 1301–1313

44. Aksu-Menges, E., Eylem, C. C., Nemutlu, E., Gizer, M., Korkusuz, P., Topaloglu, H., Talim, B., and Balci-Hayta, B. (2021) Reduced mitochondrial fission and impaired energy metabolism in human primary skeletal muscle cells of Megaconial Congenital Muscular Dystrophy. Sci Rep 11, 18161

45. Tezze, C., Romanello, V., Desbats, M. A., Fadini, G. P., Albiero, M., Favaro, G., Ciciliot, S., Soriano, M. E., Morbidoni, V., Cerqua, C., Loefler, S., Kern, H., Franceschi, C., Salvioli, S., Conte, M., Blaauw, B., Zampieri, S., Salviati, L., Scorrano, L., and Sandri, M. (2017) Age-Associated Loss of OPA1 in Muscle Impacts Muscle Mass, Metabolic Homeostasis, Systemic Inflammation, and Epithelial Senescence. Cell Metabolism 25, 1374–1389.e6

46. Buck, M. D., O’Sullivan, D., Klein Geltink, R. I., Curtis, J. D., Chang, C.-H., Sanin, D. E., Qiu, J., Kretz, O., Braas, D., van der Windt, G. J. W., Chen, Q., Huang, S. C.-C., O’Neill, C. M., Edelson, B. T., Pearce, E. J., Sesaki, H., Huber, T. B., Rambold, A. S., and Pearce, E. L. (2016) Mitochondrial Dynamics Controls T Cell Fate through Metabolic Programming. Cell 166, 63–76

47. D’Amico, D., Mottis, A., Potenza, F., Sorrentino, V., Li, H., Romani, M., Lemos, V., Schoonjans, K., Zamboni, N., Knott, G., Schneider, B. L., and Auwerx, J. (2019) The RNA-Binding Protein PUM2 Impairs Mitochondrial Dynamics and Mitophagy During Aging. Mol Cell 73, 775–787.e10

48. Popov, V., Medvedev, N. I., Davies, H. A., and Stewart, M. G. (2005) Mitochondria form a filamentous reticular network in hippocampal dendrites but are present as discrete bodies in axons: a three-dimensional ultrastructural study. J Comp Neurol 492, 50–65

49. Picard, M., White, K., and Turnbull, D. (2012) Mitochondrial morphology, topology, and membrane interactions in skeletal muscle: A quantitative three-dimensional electron microscopy study. Journal of applied physiology (Bethesda, Md. : 1985) 114

50. Iqbal, S., Ostojic, O., Singh, K., Joseph, A.-M., and Hood, D. A. (2013) Expression of mitochondrial fission and fusion regulatory proteins in skeletal muscle during chronic use and disuse. Muscle & Nerve 48, 963–970

51. Zhang, L., Trushin, S., Christensen, T. A., Bachmeier, B. V., Gateno, B., Schroeder, A., Yao, J., Itoh, K., Sesaki, H., Poon, W. W., Gylys, K. H., Patterson, E. R., Parisi, J. E., Diaz Brinton, R., Salisbury, J. L., and Trushina, E. (2016) Altered brain energetics induces mitochondrial fission arrest in Alzheimer’s Disease. Sci Rep 6, 18725

52. Mishra, P. and Chan, D. C. (2016) Metabolic regulation of mitochondrial dynamics. Journal of Cell Biology 212, 379–387

53. Fuqua, J. D., Lawrence, M. M., Hettinger, Z. R., Borowik, A. K., Brecheen, P. L., Szczygiel, M. M., Abbott, C. B., Peelor, F. F., Confides, A. L., Kinter, M., Bodine, S. C., Dupont-Versteegden, E. E., and Miller, B. F. (2023) Impaired proteostatic mechanisms other than decreased protein synthesis limit old skeletal muscle recovery after disuse atrophy. J Cachexia Sarcopenia Muscle

54. Augusto, V., Padovani, C., Eduardo, G., and Campos, R. (2004) Skeletal muscle fiber types in C57BL6J mice. J. morphol. Sci 21, 89–94

55. Joseph, A.-M., Adhihetty, P. J., Buford, T. W., Wohlgemuth, S. E., Lees, H. A., Nguyen, L. M.-D., Aranda, J. M., Sandesara, B. D., Pahor, M., Manini, T. M., Marzetti, E., and Leeuwenburgh, C. (2012) The impact of aging on mitochondrial function and biogenesis pathways in skeletal muscle of sedentary high- and low-functioning elderly individuals. Aging Cell 11, 801–809

56. Leduc-Gaudet, J.-P., Picard, M., Pelletier, F. S.-J., Sgarioto, N., Auger, M.-J., Vallée, J., Robitaille, R., St-Pierre, D. H., and Gouspillou, G. (2015) Mitochondrial morphology is altered in atrophied skeletal muscle of aged mice. Oncotarget 6, 17923–17937

57. Ferreira, R., Vitorino, R., Alves, R. M. P., Appell, H. J., Powers, S. K., Duarte, J. A., and Amado, F. (2010) Subsarcolemmal and intermyofibrillar mitochondria proteome differences disclose functional specializations in skeletal muscle. PROTEOMICS 10, 3142–3154

58. Balan, E., Schwalm, C., Naslain, D., Nielens, H., Francaux, M., and Deldicque, L. (2019) Regular Endurance Exercise Promotes Fission, Mitophagy, and Oxidative Phosphorylation in Human Skeletal Muscle Independently of Age. Front. Physiol. 10

59. Jiang, X. L., Gu, X. Y., Zhou, X. X., Chen, X. M., Zhang, X., Yang, Y. T., Qin, Y., Shen, L., Yu, W. F., and Su, D. S. (2019) Intestinal dysbacteriosis mediates the reference memory deficit induced by anaesthesia/surgery in aged mice. *Brain*, Behavior, and Immunity 80, 605–615

60. Guo, L. Y., Kaustov, L., Brenna, C. T. A., Patel, V., Zhang, C., Choi, S., Halpern, S., Wang, D.-S., and Orser, B. A. (2023) Cognitive deficits after general anaesthesia in animal models: a scoping review. British Journal of Anaesthesia 130, e351–e360

